# Re-invasion of the New World screwworm in Central and North America: comparative analysis of three SIT eradication efforts

**DOI:** 10.64898/2026.07.16.738939

**Authors:** Andrew Paul Gutierrez, Luigi Ponti

## Abstract

In sharp contrast to biological control that seeks control of pests below economic levels, eradication programs seek total extermination of pest species. However, eradication programs are often undertaken without knowledge of the pest’s potential geographic distribution and relative abundance (i.e., geographic risk assessment), often leading to eradication efforts conducted across areas much greater than the potential permanent range of the target species. To demonstrate this, we review eradication efforts against the tropical New World screwworm (*Cochliomyia hominivorax*) in the SE USA and Mexico, the exotic sub-tropical tephritid Mediterranean fruit fly (*Ceratitis capitata,* medfly) in California, and the exotic tropical pink bollworm (*Pectinophora gossypiella*, PBW) in the SW USA. We use mechanistic physiologically based demographic models (PBDMs) that are physiologically based time varying life tables to decompose the systems and to assess risk under extant and climate change scenarios.

## Introduction

Globally, invasive species from a wide variety of taxa are extending their range and/or invading new areas with unknown consequences in the face of extant weather and global and climate change. For an invasive species to successfully invade new areas, its life history traits and biology (physiology and behavior) must enable them to establish and survive time-varying patterns of weather (temperature, relative humidity, and other factors), and biotic interactions (available hosts, competitors, natural enemies) (e.g., Gutierrez 1996, Myers et al. 2000, Sequeira et al. 2001). Recognition of such constraints has long been part of ecological theory (Andrewartha and Birch 1954) and biological control (e.g., Huffaker and Messenger 1976). Furthermore, the risk from target pest species must be assessed under extant and climate change weather, and yet the question of how to analyze the inherent complexity of invasive species problems as driven by weather in natural and agricultural ecosystems is thought to be unresolved (Dormann et al. 2017).

Some exotic species establish and appear to cause little harm to the environment or agriculture, but others cause serious damage and must be controlled or managed below economic levels using various methods, including eradication (see Ricciardi et al. 2021). Some invasive species are not amenable to biological control because no effective natural enemies exist and/or because the *ex-ante* prerelease efficacy of the control agents is unknown (Gutierrez, Ponti, Neuenschwander, et al. 2025). Such species are often targets for eradication using the sterile insect technique (SIT; Knipling 1955) the simple theoretical underpinning of which are: (1) the target population has a ‘stable’ population closed to immigration, (2) uniform age structure, (3) discrete generations, (4) a constant *per capita* reproductive rate, and (5) the release of sterile males must be high enough to achieve insemination of virgin wild females by sterile SIT males despite competition from wild, fertile males (*cf.* Krafsur 1994). We note that conditions 1-4 are unlikely to be met in the field.

Government-sponsored eradication programs using a wide variety of techniques have collectively had a checkered history of success. Among these methods are meiotic drive, mass trapping, lethal or sterility genes, *Wolbachia* infection that hinders reproduction (incompatible insect technique, IIT), area-wide suppression including mass trapping and mating disruption, microbial insecticide sprays, transgenic crops (e.g., Bt crops), mechanical control, barrier zones to slowing the spread, the sterile insect technique (SIT), and others (see e.g., Myers et al. 2000, Martinez et al. 2020). Pérez-Staples *et al*. (2021) reviewed the SIT eradication literature for the Neotropics. What is clear is that the ecological bases for most eradication programs have been highly suspect, with eradication efforts often claiming success in marginal areas where the target species cannot survive (e.g., tropical fruit flies in temperate areas; Gutierrez et al. 2021). In such cases, eradication programs adopt an old axiom of war: “*declare victory (even in defeat) and leave*”!

A basic question that should be answered before launching eradication efforts: What are the biotic and abiotic factors that determine the prospective geographic range and relative abundance of the target species? Sadly, this is seldom addressed with clarity, and verbal non-quantitative assessments (Carey and Dowell 1989), statistical inference of detection data (Papadopoulos et al. 2013), and various correlative climate-based ecological niche models (ENMs, e.g., CLIMEX, MaxEnt) are invoked; methods that tell us little about the biology of invasion or the potential for control. ENMs require the use of species occurrence data as a basis for determining the ecological correlates in rough aggregate weather data responsible for the observed distribution and the prospective distribution under extant and climate change conditions. The shortcomings of ENMs have been widely reviewed (e.g., Fischlin et al. 2007, Elith and Leathwick 2009, IPCC 2014).

Addressing the above questions requires “… *models based on understanding the processes that result in a system behaving the way it does, … remaining valid indefinitely*” (Evans 2012). We must seek reliable rules (”Laws”) of nature common to all species and ecosystems that, in the best case, include thermodynamics and mathematical descriptions of population and trophic interactions (Jørgensen et al. 2016). This is an exceedingly high standard, and as an intermediate step, we propose mechanistic physiologically based demographic models (PBDMs, i.e., physiologically based time varying life tables) based on simple bioeconomic rules of resource acquisition and allocation common to all species (e.g., Gutierrez 1996, Ponti et al. 2019). [We apologize for the unavoidable self-reference as many of the PBDM were developed by us and research associates.]

PBDMs capture the weather-driven biology of species and, in a tri-trophic context, biotic interactions, and the models are transferable across time and place (Gutierrez and Ponti 2013). Although PBDMs are not couched in terms of physics, they do conform to the first two Laws of thermodynamics. The underlying bases and mathematics of the PBDM paradigm are outlined in Gutierrez (1996) and Ponti *et al*. (2019). In a bio-economic context, PBDMs may be developed by modeling the processes of energy acquisition and allocation (metabolic pool models, MPMs; see Gutierrez 1992, 1996). Another approach is to parameterize the model using biodemographic functions (BDFs) derived from age-specific life table studies across abiotic gradients (Gutierrez, Ponti, Neteler, et al. 2025). In the BDF approach, the estimated functions are the results of resource acquisition and allocation under the experimental conditions. In both cases (MP and BDF), the models are resource demand-driven and supply/demand ratio-dependent, and both are dynamic weather-driven time-varying life tables (*sensu* Gilbert et al. 1976, Gutierrez 1996). In some cases, the PBDM may be constructed as hybrid of the two approaches – it all depends on the available data and the question to be answered. The theoretical basis of PBDMs was reviewed by Gutierrez (1992) and Mills and Gutierrez (1999). Barlow (1999)(1999) supposed that PBDMs have large numbers of parameters and are difficult to develop, but this is not the case, as demonstrated by numerous studies summarized in the supplemental materials in Gutierrez et al. (2025) where the same underlying model was used (e.g., Gutierrez and Ponti 2013). Here, three species we have studied intensively are used to demonstrate the PBDM analysis approach in the evaluation of the SIT technology.

This paper extends the analyses of the invasive potential and eradication efforts of the tropical new world screwworm (*Cochliomyia hominivorax*) in the SE USA and Mexico (Gutierrez and Ponti 2013, 2014, Gutierrez et al. 2019), the exotic subtropical tephritid Mediterranean fruit fly (*Ceratitis capitata,* medfly) in California (Gutierrez and Ponti 2011), and the exotic subtropical pink bollworm (*Pectinophora gossypiella*, PBW) in the SW USA (Gutierrez et al. 2006). None of the three species has effective natural enemies, and eradication was attempted despite incomplete understanding of their weather-related ecology and biology and of an analysis of their potential geographic distribution (Gutierrez and Ponti 2022). Also reviewed here are their prospective changes in the geographic range under climate change.

### Eradication of New World screwworm – an ongoing 60-year effort

The new world screwworm fly is a subtropical-tropical species of the Americas that periodically caused outbreaks of myiasis in the SW USA, primarily south and central Texas, and less so in neighboring states (OIE 2013). The SIT-based eradication of the fly from North America to the Darian Gap in Panama during 1957-2001 was heralded as the quintessential success of the sterile insect technology (SIT) proposed by Knipling (1955). The SIT model provided the principles (see above) upon which the USDA and the Mexico-US Commission planned the screwworm eradication operations using releases of prodigiously large numbers of irradiated sterile male flies targeting unmated adult females, chemical treatment of infested livestock to kill larval stages in wounds and adults feeding on serous fluid in wounds, and the establishment of quarantine areas to prevent the introduction of infested animals. Valdez-Espinoza et al. (2025) review the potential path of the reinvasion of Mexico from Central America.

Surprisingly, despite nearly a billion dollars spent on screwworm eradication, studies of its thermal biology of development, reproduction, and mortality are sparse in the literature, greatly hindering model development. Furthermore, although nearly 40 years have elapsed since the eradication program ended in the USA, the extensive SIT program records of field myiasis in the USA had not been tabulated nor fully analyzed (Gutierrez et al. 2019). Only Krafsur (1987) attempted to find weather correlates for screwworm outbreaks in Texas. The data for myiasis in Texas during the SIT eradication period are available at https://doi.org/10.5281/zenodo.19893675 (Gutierrez et al. 2026).

#### Biology and ecology

The model for screwworm is a PBDM/BDF model parameterized using sparse data from the literature. Extensive field studies on its ecology reported that adult male flies feed at flowers and live 2–3 weeks, while adult females live on average about 10 days, feeding on serous fluids at animal wounds and on decomposing animals (Thomas and Mangan 1989). Promiscuous mating in males (polygyny) and single mating in females is the rule and was a key factor in the SIT program. About 3-4 days (*d*) after mating, female flies begin to seek wounds on vertebrates to lay large batches of eggs. Screwworm females are attracted to wounds as small as those caused by the feeding of the invasive cattle tick (*Rhipicephalus (Boophilus) microplus*) (OIE 2013) that also has periodic outbreaks in Mexico and south Texas (Pérez de León et al. 2012). Feeding by screwworm larvae expands the wound (myiasis), attracting further oviposition, and if not treated, the myiasis may cause the death of the animal. The egg and larval stages develop at host body temperature, and at maturity, the larvae drop to the ground to pupate. Pupae and free-living adults experience near ambient temperatures. The autogenous females can complete 2-3 vitellogenic cycles without a protein meal (Crystal 1966). Under field conditions in Central America, the mean age of females at wounds was 7.5 *d* with a longevity of 21*d* (Thomas and Chen 1990). Screwworm is cold-intolerant and has high lower and upper developmental thermal thresholds (14.5 and 43.5 °C, respectively) with the optimal temperature for survival and adult reproduction being approximately 27.5 °C (partial data from Adams 1979, Berkebile et al. 2006, Gutierrez et al. 2019). The fly lacks the capacity for diapause, further restricting its northern range in North America.

#### Screwworm invasion biology

The following summary of the invasion biology of screwworm into the SE USA is based on a review of the literature and PBDM analyses by Gutierrez and Ponti (2014) and Gutierrez *et al*. (2019). Cold temperatures limit the potential northward endemic range of the fly by limiting reproduction and increasing winter mortality. The predicted northernmost marginal limit of potential permanence in the USA is the tip of South Texas and Florida, but this may change with climate warming. The analysis found that outbreaks of myiasis in Texas are a two-year process with screwworm adults migrating northward from the warmer southern endemic areas of Mexico on wet North American monsoon winds. For this to occur, conditions in the transition area of NE Mexico and south Texas have to have favorable warmer winters the previous year (*y-1*) allowing population survival and population buildup in early spring. Aided by southerly monsoon winds (northward flow), the fly disperses into Texas during the spring-summer of year *y,* causing myiasis outbreaks across Texas and elsewhere. This bi-seasonal invasion biology explained the series of outbreaks that occurred in Texas during the 1962-82 period when the eradication efforts were in full force (Gutierrez et al. 2019).

Prevention of screwworm reinvasion of Texas required eradication of the fly in endemic subtropical and tropical areas of Mexico and Central America. Field studies showed that despite screwworm’s high reproductive potential, conditions that enabled its eradication in the tropics include low growth rates of endemic field populations (Thomas and Mangan 1992) with field doubling times ranging from 54 to 139 days (Matlock and Skoda 2009). Further, oviposition site densities (wounds) are generally low, and the innate success rate in finding them is low, resulting in boom-to-bust reproductive dynamics as oviposition sites and weather allow (Krafsur et al. 1980, Krafsur 1998). In the subtropics, a boom phase may occur when weather is warm and oviposition sites are available, while a bust phase occurs as modest declining temperatures decrease fly vital rates and increase mortality rates. The added load of massive sterile fly releases can drive intrinsically low screwworm populations to demographic mate-limited “Allee” extinction (*cf.* Courchamp et al. 2008, Gutierrez et al. 2019). In tropical areas such as Tuxtla-Gutierrez, Mexico, and Panama, cool weather effects are weak, and high levels of SIT releases are required for eradication. Furthermore, polygyny in males and single mating in females make screwworm highly susceptible to massive SIT releases, but even then, SIT releases appear to have a low efficacy rate (<1.25%; Krafsur 1985, Gutierrez et al. 2019). Hence, the gradients of low climatic favorability in most of Texas to high favorability in tropical areas of southern Mexico and Central America required different SIT release rates for eradication.

#### SIT eradication efforts

SIT efforts began in Florida in 1957, in Texas in 1962, and progressed through Mexico during the 1980s to the Darien Gap in South Panama in the late 1990s (Wyss 2000), where containment efforts continue to keep the fly from reinvading northward (Maxwell et al. 2017). During the 1957 - 2000 period, eradication of the screwworm cost more than 750 million US dollars, with ongoing annual costs of >$15 million to produce millions of sterile flies in the Panama production facility for containment releases over eastern Panama and areas of Colombia (USDA-APHIS 2017a). The containment efforts in Panama continue as the observed “…increased number of (myiasis) cases in […] clusters could be due to SIT failure, the regular transport of screwworm-positive animals …, movement of screwworm-positive wildlife and a lack of fly control in neighboring Colombia” (Maxwell et al. 2017, see Franco-Molina et al. 2025, Zaldivar-Gomez et al. 2025). The fly has now escaped containment in Panama and reinvaded Central America and parts of Mexico, as documented by Valdez-Espinoza et al. (2025). Furthermore, screwworm is endemic to the Caribbean and South America, and periodic cases of myiasis occur in North America (Alexander 2006), as exemplified by the severe outbreak of myiasis in deer (*Odocoileus virginianus clavium*) in the Florida Keys in 2016. This infestation was eradicated by releasing 188 million sterile flies (USDA-APHIS 2017b, Skoda et al. 2018).

Such problems are harbingers of difficulties that may be encountered if attempts are made to extend eradication across the vast tropical and subtropical areas of South America (Gutierrez et al. 2019).

#### Climatic limits for screwworm in North America

The prospective permanent range of the tropical screwworm is determined by cold weather as measured by the annual sum of daily mortality rates for pupae and adults computed daily (*d*) for each of 15,000 lattice cells in the USA and Mexico 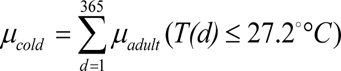). The average annual sum across years for extant weather (*n* = 1975 − 2005), and for climate change weather (*n* = 2045 − 2055) were computed as 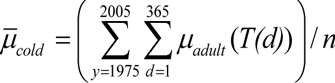. The values truncated to 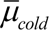 ≤ 10 are summarized in Figure 1a,b (from Gutierrez et al. 2019). The simulation results suggest that the average zone of permanent habitation is 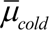 2.5, though the actual geographic range of permanence may vary annually as weather patterns change. Gutierrez and Ponti (2014) estimated that an average value of 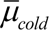≈ 10 was a good metric defining the geographical limits for screwworm that accorded with field observations, but the limit of endemicity is better defined by 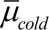 an obvious error in Gutierrez and Ponti (2014).

**Figure 1.**
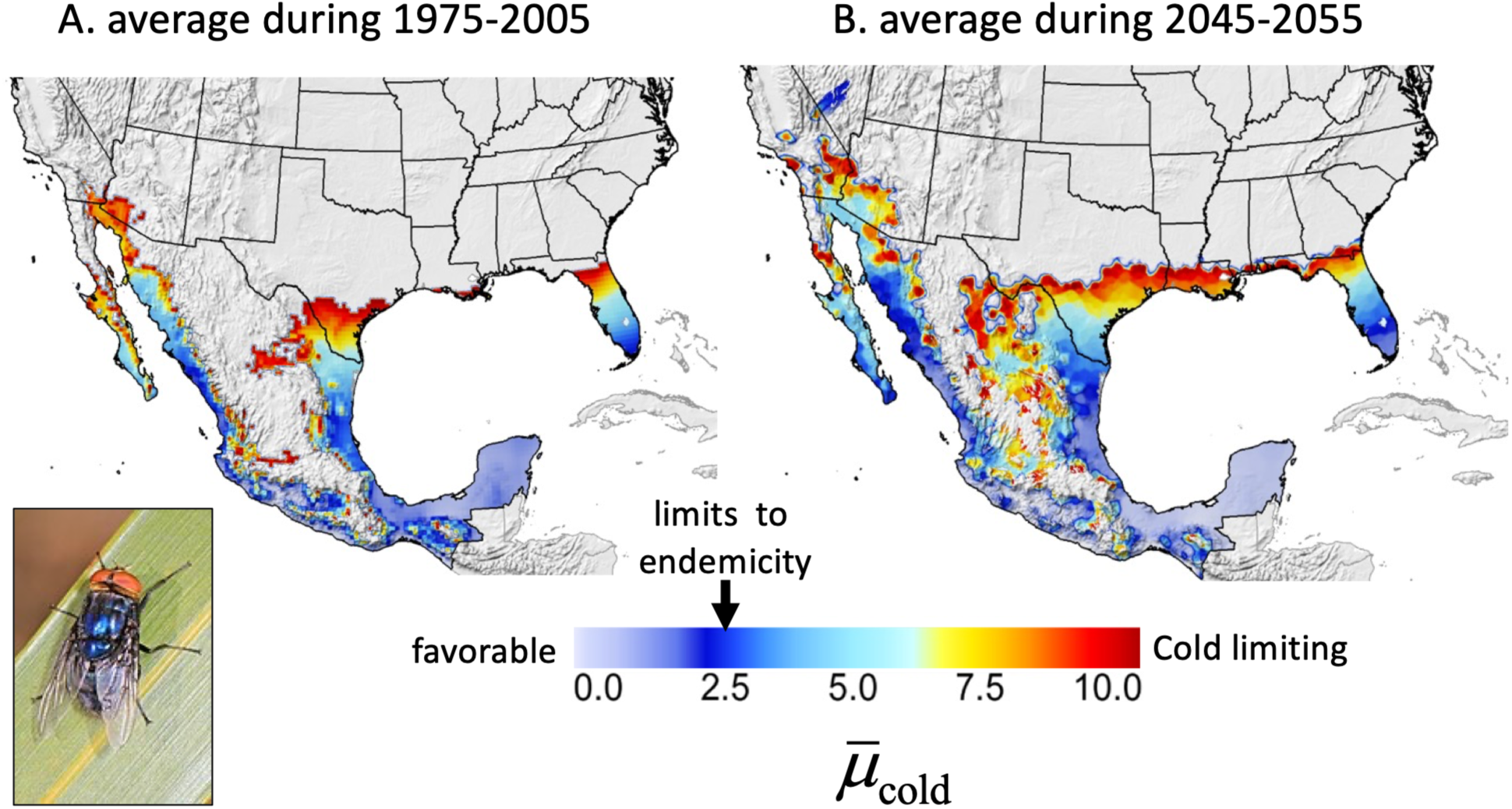
Comparison of areas favorable for screwworm endemicity in the USA and Mexico (excluding the Caribbean and Central America where it was endemic) based on annual cumulative average cold-temperature mortality rates (μ_cold_) computed from daily weather data from a high resolution and bias-corrected climate scenario (Thrasher et al. 2012) for: (a) the historical period 1975–2005 and (b) the future period 2045–2055 (from Gutierrez et al. 2019). High values of μ_cold_ (dark red) and the large grey areas beyond the margins of μ_cold_ =10 being unfavorable, whereas light blue-grey simulated areas are increasingly favorable in the range 0 ≤ μ_cold_ *<*2.5. Inset for screwworm adult is from https://commons.wikimedia.org/wiki/File:Screwworm_-_Cochliomyia_hominivorax,_Big_Pine_Key,_Florida.jpg

#### Climate change effects

Using high resolution, bias-corrected NASA climate model data for 2045-2055 (see Thrasher et al. 2012, NASA 2015), the PBDM predicts increases in the fly’s potential endemic range into the southern USA and higher elevations in Mexico. The prospective distribution is relative due to the difficulties of predicting future weather. However, an increase of 2° C (or more) in average temperatures by the year 2050 would increase the prospective limits of screwworm endemicity into the southern reaches of Texas and Florida (μ_cold_ *<*2.5; Fig. 1a vs Fig. 1b) (Gutierrez et al. 2019). Under climate change, the prospective areas of endemicity in Mexico and southern Texas and Florida would increase, and hence the area of the USA invaded due to the monsoon winds would increase, challenging the low efficacy SIT interventions as occurred in Texas during the large 1972 outbreak, and the predicted potential outbreaks predicted during 1982 – 2010 (see Gutierrez et al. 2019).

However, SIT eradication of screwworm up to the Darian Gap in Panama was successful (Krafsur et al. 1986, Comis et al. 2012), but extensive eradication efforts in the USA occurred largely outside areas of the pest’s climate envelope of endemicity (e.g., most of Texas). This contrasts with the outbreak that occurred in 2016 in the Florida Keyes where endemicity is likely.

### Eradication of Mediterranean fruit fly – a questionable success

The sub-tropical medfly is a polyphagous fruit pest originally from sub-Saharan Africa and is a serious pest in many tropical regions of the world. Medfly is established in near coastal regions of the Mediterranean Basin. In the western hemisphere, it was first reported in Costa Rica in 1955, invaded progressively northward, and could infest all Central American countries (Vail et al. 1976) and tropical areas of South America. We focus on North America, where extensive eradication efforts on medfly occurred. Medfly was eradicated from Mexico and Guatemala, where ongoing prophylactic eradication efforts continue. The literature on medfly biology is large, consisting of panoply studies on a wide variety of topics, but provides sufficient data to parameterize the PBDM (Gutierrez and Ponti 2011, Gutierrez et al. 2021). Detection records for medfly are available as supplemental material in Papadopoulos *et al*. (2013), that we argue are for transient infestations (Gutierrez et al. 2014).

The potential threat from tropical tephritid fruit flies has been couched in economic terms of a multi-billion-dollar threat to agriculture (Carey et al. 2017), and a substantial surveillance and eradication infrastructure has developed in response. On average, an eradication campaign against fruit flies in California is estimated to cost approximately US$32 million (Papadopoulos et al. 2013) and up to US$100 million as occurred for medfly in the San Francisco Bay Area in 1980-81 (Carey 2010). For example, the average cumulative eradication cost for oriental fruit fly (*Bactrocera dorsalis*) in 166 lattice cells of size 100 km^2^ in California from 1960 through 2017 was US$610 million (Zhao et al. 2019). Government eradication agencies claim that incipient infestations of tropical fruit flies have been eradicated using combinations of insecticides and SIT, and yet they fail to explain why other species such as the olive fly (*Bactrocera oleae*) (Gutierrez et al. 2009) and spotted wing drosophila (*Drosophila suzukii*) (see Gutierrez et al. 2016) have established widely in California, or why no attempt to eradicate them has occurred (Gutierrez et al. 2021).

#### Eradication efforts

Medfly was posited eradicated from Mexico but reinfestations likely occur. The fly is established in Hawaii, and has been detected primarily in Florida, and in the coastal plain from Los Angeles to San Diego, but also in the south San Francisco Bay area of California (see Papadopoulos et al. 2013). Infestations in California during the late 1970s and early 1980s garnered considerable political and applied scientific attention. Carey and Dowell (1989) and Carey *et al*. (2017) gave a verbal rationale for medfly’s invasive potential in California, arguing that it has climatic analogs to the Mediterranean basin, where medfly is established in some areas. Carey (1996) proposed a predictive framework for medfly invasion of California, including maps of invasion pathways. More recently, Papadopoulos *et al*. (2013) claimed that at least five to nine invasive fruit fly species (including medfly) are established in California, but at undetectable levels – a claim that cannot be verified nor falsified – a claim made without analyses of the role of weather in their potential for establishment and geographic range (see Gutierrez and Ponti 2011, Gutierrez et al. 2014, McInnis et al. 2017). Carey *et al*. (2017) defended these claims, ignoring the argument of McInnis *et al*. (2017) and Gutierrez et al. (2014) that if the tropical fruit flies are established in California and the climate is favorable, then fly population growth would occur and the flies would be detectable. Prognostications by Carey and colleagues also ignored microsatellite and mitochondrial DNA evidence of multiple introductions of medfly (e.g., Davies et al. 1999, Meixner et al. 2002, see Liebhold et al. 2010), and dismissed as faulty the results of a weather-driven PBDM analysis for medfly that prospectively showed that most of California was unfavorable, and that only the coastal areas of southern California were marginally favorable (Fig. 5a and b in Gutierrez and Ponti 2011). And yet, the same model accurately predicted the distribution in the European-Mediterranean region in areas where establishment is detected (see Gutierrez et al. 2021). Medfly is not considered established (Dr. Kyle Beucke, Primary State Entomologist, California Department of Food and Agriculture (CDFA), personal communication), but establishment may change with climate warming (Gutierrez et al. 2021).

#### Prospective range under extant and climate change weather

Figure 2a shows the prospective area of favorability in North America and as an inset in California under extant 1980-90 weather, while Figure 2b and inset show the changed prospective range under projected 2060-65 climate change weather. Comparing the two scenarios is difficult because under 1980-90 weather, in areas of Mexico and Central America, maximum favorability (FI∼1) occurs with predicted cumulative 8,500 pupae per year, while in marginal coastal southern California with FI≤0.5, ∼3,200 pupae are predicted. Under climate change, the numerical scale doubles, the prospective favorability in coastal southern California as measured by pupal density increases >4fold with fly densities greater than those predicted for tropical Mexico – Central America during the 1980-1990 period when the fly was known capable of infesting those areas (Gutierrez et al. 2021).

**Figure 2.**
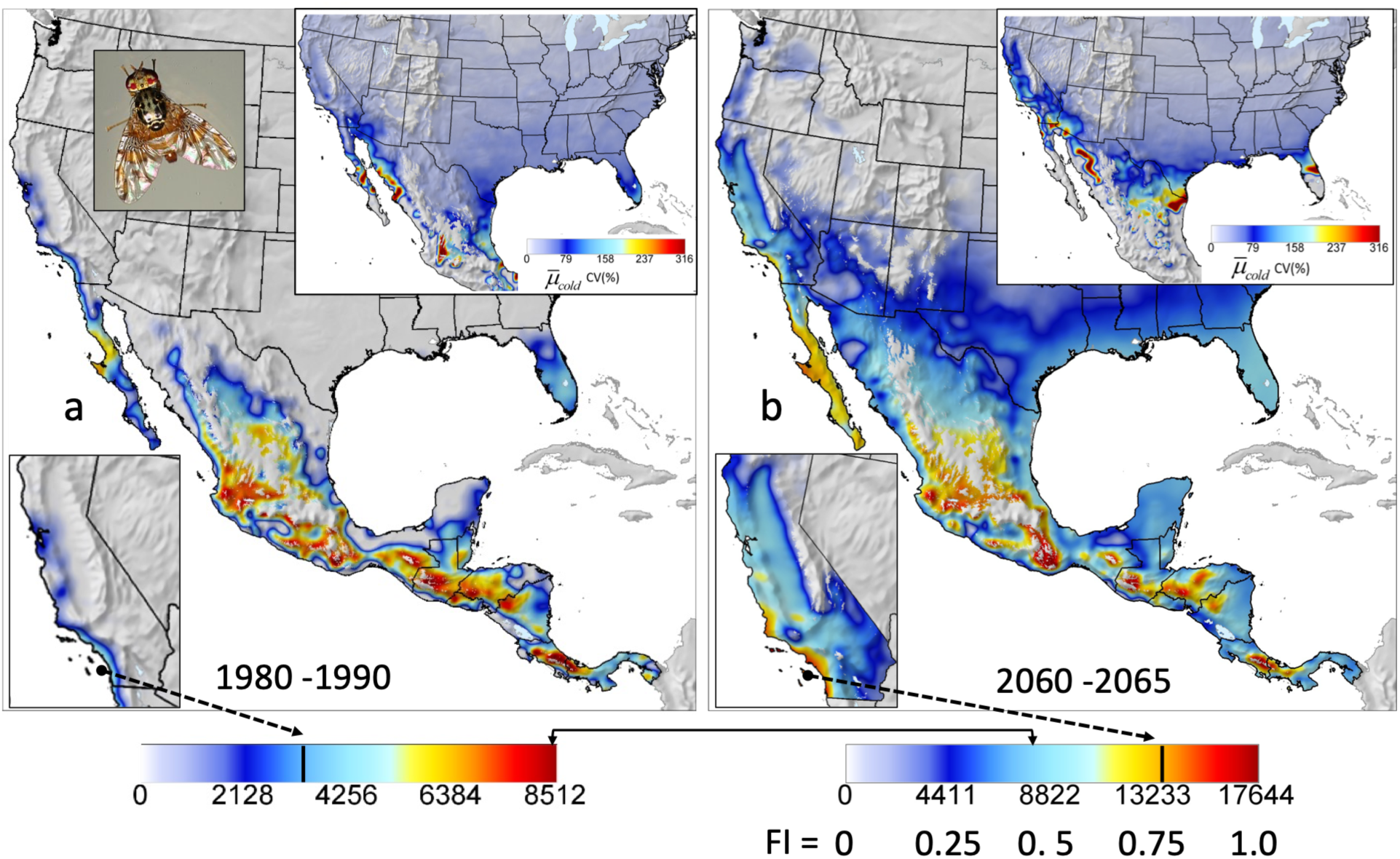
Simulation of medfly distribution and relative abundance in North and Central America below 2000m: (a) during 1980-1990, and (b) under climate change (2060-2065). Inset for medfly adult female is from https://commons.wikimedia.org/wiki/File:Mediterranean_Fruit_Fly_Female_(8543655978).jpg

This suggests that under climate change, medfly could be established in coastal California and in few inland locations. A measure of the favorability of temperature in each lattice cell is the annual sum of daily mortality rates below the lower thermal threshold that occur during the year (i.e., 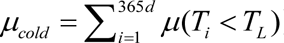), with means and coefficients of variation computed from the annual values (see Gutierrez et al. 2021). Insets in Figures 2a and 2b (upper right) show a band of high variability of 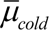 (CV(%)∼80) in a transition zone between northern areas of inhospitability (i.e., temperate areas with consistently cold limiting weather with low CV) and southern regions of increasing hospitability (i.e., consistently favorable weather with low CV). For 1980-90 weather, this zone is further south and weakly defined, while for future climate, this transition band stretches from northern California into northern Mexico and across Texas and northern Florida. Under climate change, the prospective range of medfly in Mexico and Central America is projected to increase (i.e., values above 8,000 in Fig. 2b).

In summary, the 1970-2010 period in California was not particularly favorable for medfly (FI<0.5; see Gutierrez and Ponti 2011, Gutierrez et al. 2021), and though not available at the time, PBDM assessment could have aided eradication agencies in their efforts to evaluate the threat medfly posed. We note, however, that with 1.5-2 °C climate warming by 2060-2065, coastal California will become more tropical, and conditions will become favorable for medfly to establish at detectable levels (lower left insets in Fig. 2a vs 2b).

### Eradication of pink bollworm – a documented success

The stenophagous pink bollworm (PBW) is a tropical species native to South Asia (Papua New Guinea–North Australia; van den Bosch and Messenger 1973, see Grefenstette et al. 2009). It is widely distributed in hot, dry cotton-growing areas worldwide, where it also attacks other species of Malvaceae (e.g., okra, hibiscus, hollyhock). PBW has no effective natural enemies. PBW was first discovered in Florida in 1932 on tree cotton and spread to commercial cotton in the US and Mexico. The moth invaded the desert cotton areas of Arizona and southern California in the late 1960s, where it became the key pest and, due to the extensive use of insecticides to control it, engendered massive outbreaks of secondary pests (e.g., bollworm, budworms, whitefly, and mites). Dispersal of PBW in California is aided by southwesterly monsoon winds that annually carry adult moths from the southern desert valleys of California northward into the southern part of the Central Valley and elsewhere (Stern and Sevacherian 1978). Light infestations were detected in the southern reaches of the Central Valley in late summer, but analysis shows the moth cannot overwinter under extant climatic conditions (Gutierrez et al. 2006).

The extensive literature on the moth’s biology and ecology was used to develop PBDMs for cotton and for PBW (see Gutierrez et al. 1977). Critical elements of pink bollworm biology include tight links to the phenology and dynamics of cotton fruiting. Important components of PBW’s biology include diapause initiation in mature larvae during late summer in response to decreasing photoperiod and temperature, the phenology of spring emergence (Gutierrez et al. 1981), and cold-intolerance of diapause larvae (Gutierrez et al. 1977, 2006, data from Venette et al. 2000). A PBDM/MP model for cotton and a PBDM/BDF model for pink bollworm were used to assess the biology, geographic distribution, and relative abundance of the PBW (Gutierrez et al. 1977, Stone and Gutierrez 1986), including the effects of highly efficacious Bt cotton (Gutierrez and Ponsard 2006).

Figure 3 shows simulated (a) prospective irrigated cotton yields and three measures of climatic favorability for PBW across the USA and Mexico under extant weather: (b) average cumulative larvae plant^-1^ *d^-1^ y^-1^*(i.e., larval days during the season), (c) average number of diapause larvae plant^-1^y^-1^, and (d) normalized average winter survival of diapause larvae. Prior to the introduction of Bt cotton, high densities of summer larval populations were common in cotton (Fig. 3b) and large number of diapause larvae developed in southern California desert valleys and in Arizona (Fig. 3c). However, winter survival is not predicted in the Central Valley of California (see Gutierrez et al. 2006), and over much of the cotton belt in the SE US and most of Texas, southern New Mexico, and north-central Mexico (Fig. 3d). High winter survival is predicted in the Yucatan Peninsula, but an adverse combination of high fall temperatures and short photoperiod reduce diapause induction there (Fig. 3c) (see Gutierrez et al. 1981). PBW is not a serious pest in Central America (https://www.cabi.org/isc/datasheet/39417) because of short daylength and seasonal planting of cotton.

**Figure 3.**
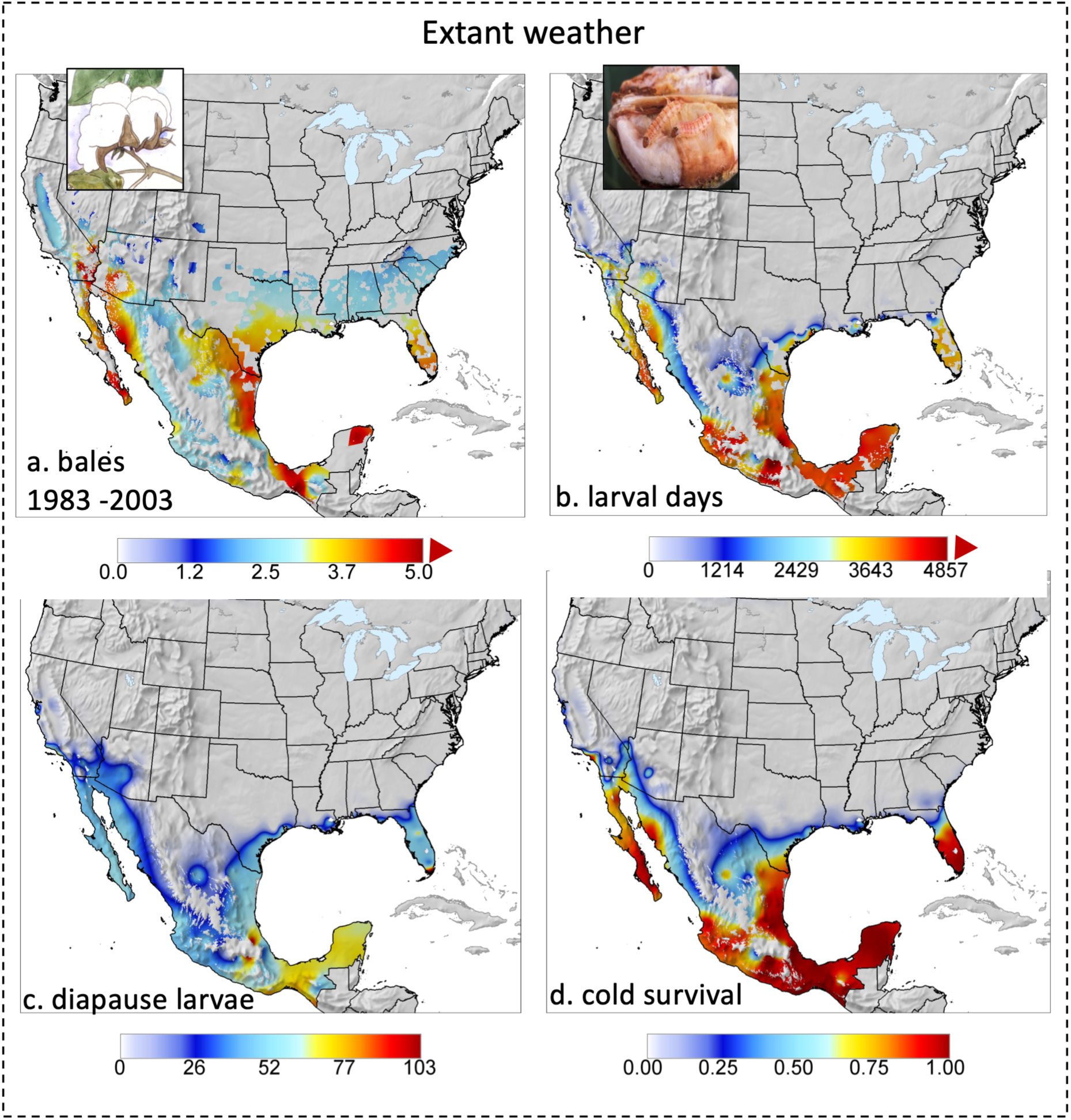
Geographic limits of pink bollworm in the SW USA during 1983-2003: (a) bales/acre (450 lbs of seed cotton per bale), (b) cumulative larval days, (c) number of diapause larvae m^-2^, and (d) index of cold weather survivorship (see text, modified from Gutierrez and Ponti 2013, 2022). Insect for pink bollworm larvae is from https://commons.wikimedia.org/wiki/File:Pinkbollworm.jpg

Under 2060-2065 climate change weather, prospective cotton yield is predicted to decrease in North America including the Great Central Valley of California (Fig. 4a), but absent eradication, the range of PBW would expand northward into the great central valley of California and decrease in Mexico (Fig. 4b). Compared to extant weather (Fig. 3), the numbers of diapause larvae would decrease (Fig. 4c vs Fig. 3c), and cold survival would decrease in some areas of Mexico but would increase in Western Mexico into California (4d vs 3d). Figures 4e and 4f compare larval days for both the 1983-2003 and 2060-2065 periods and show the expansion of PBW into central California and in Western Mexico.

**Figure 4.**
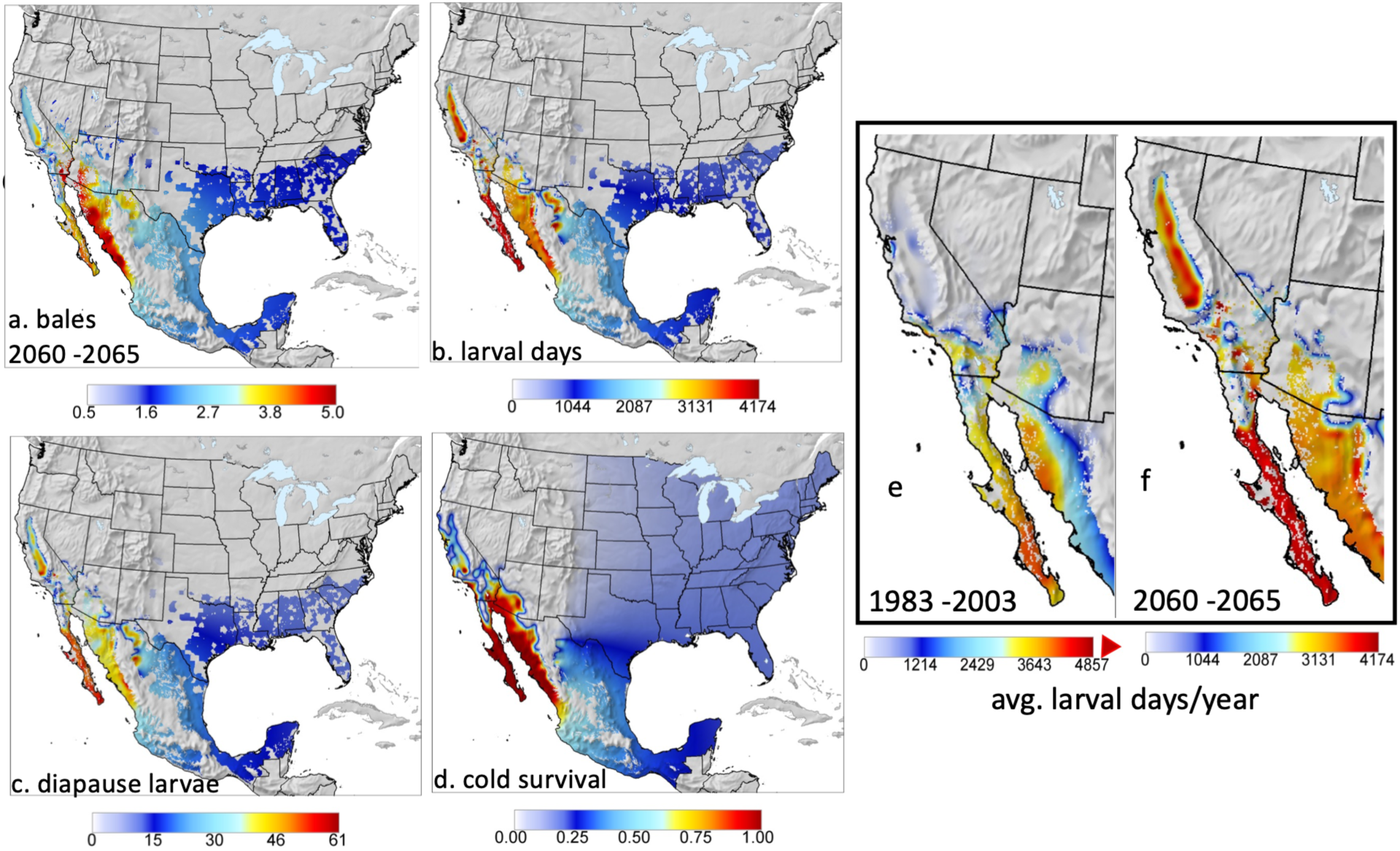
Geographic limits of pink bollworm in North America during 2060-2065: (a) bales/acre, (b) cumulative larvae days, (c) the number of diapause larvae m^-2^, and (d) cold weather survivorship (see text, modified from Gutierrez and Ponti 2013, 2022). Subfigures 4e and 4f compare average larval days per year during 1983-2003 and 2060-2065 climate warming.

#### Eradication of PBW

In 1968, the USDA and the California Department of Agriculture (CDFA) began an eradication program in Arizona and California using SIT (Staten et al. 1992), that proved elusive, and the program was redirected to PBW management using short season high density cottons and early crop termination and plowing of crop stubble that prevented the development of overwintering diapause larvae (Chu et al. 1996). A major goal was preventing the posited establishment of pink bollworm in Central Valley cotton. In 1997, genetically modified Bt cotton expressing the protoxin of the bacterium *Bacillus thuringiensis* (Bt) was introduced in southern CA and AZ (but not in the Central Valley of California), replacing the short-season cotton technology (Godfrey 2004, Gutierrez et al. 2006). Bt cotton proved highly effective against pink bollworm and greatly reduced population levels (e.g., Tabashnik et al. 2010), as it was easy to implement and reduced insecticide use and associated secondary pest outbreaks. The resultant low PBW densities achieved using Bt cotton provided conditions for renewing SIT eradication efforts.

In 2001/2002, a three-phase SIT eradication program piggybacked on the Bt cotton technology was initiated in the USA and Mexico (Grefenstette et al. 2009). Phase 1 targeted the El Paso/Trans Pecos region of west Texas, south-central New Mexico, and northern Chihuahua, Mexico – areas marginal for PBW (Fig. 3c). Phase 2 was begun in Arizona and New Mexico in 2006, and Phase 3 was begun in 2007– 2008 along the Colorado River and in the desert valleys of AZ, CA and Mexico (Grefenstette et al. 2009). The Central Valley of CA was considered an area of PBW suppression and control because the USDA claimed its SIT program in the desert areas of southern California had prevented the moth’s establishment there (Staten et al. 1992). In October 2015, PBW was declared eradicated from the southwest (https://tucson.com/news/local/how-arizona-scientists-and-farmers-banished-the-pink-bollworm-from-the-southwest/article_8c9da4ff-883c-5955-9791-363dbe95ed22.html) and in October 2018, from the continental United States (https://www.aphis.usda.gov/sites/default/files/usda-pink-bollworm-proclamation.pdf) (see Tabashnik et al. 2021).

In summary, the PBDM for cotton/PBW predicted and explained the geographic distribution, dynamics, and abundance of PBW as driven by weather across the cotton-growing region of western North America. The potential geographic distribution of PBW is restricted to the areas of cotton cultivation with mild winter temperatures. The predictions of the model contrasted with the findings of Venette *et al*. (2000) that abiotic factors did not preclude PBW’s establishment over much of the cotton belt, and that its absence in much of the area was the result of federal monitoring, quarantine, and local eradication programs. The predictions of the PBDM also conflicted with the claim that the USDA SIT eradication-suppression program kept PBW from establishing in the Central Valley of California (Staten et al. 1992) under current climate. However, the combined Bt cotton-SIT eradication did eradicate PBW in the SW USA. Furthermore, the model predicts that climate warming will increase the potential range of PBW into Central California cotton, and should reinvasion occur (see Fig. 4e vs Fig. 4f), the system dynamics must be reexamined, and the model could prove useful in evaluating strategies.

## Discussion

Native and exotic pest species in many taxa may be extremely harmful in agricultural, veterinary, and human health systems, and keeping their densities below economic levels using environmentally safe methods is an important goal. This may be achieved in a variety of ways, with eradication being the most extreme and secure result when successful. Biological control, behavioral (pheromones), and cultural methods are viable alternatives for many species, while other species prove refractory and must be controlled using cultural, toxic chemical, and biotechnology approaches. The number of examples and variants of eradication/control efforts of pest species is large, but in each case, extensive knowledge of the biology and ecology of the target pest is essential for understanding the bases of the problem and for developing environmentally friendly solutions.

Eradication agencies have historically not had the capacity to assess fully the threat posed by invasive species and the factors limiting their prospective geographic range and relative abundance, as shown by our three examples. They have been unable to distinguish between areas with high and low probability of permanence and risk, and hence where and if to concentrate eradication efforts. Widely used species distribution models provide only part of the solution, as the problem must be evaluated in a time-varying dynamic manner. Otherwise, as Krafsur (1994) surmised, during eradication efforts, “… *agencies seem usually to rely on experience and overkill to achieve goals*”. But as humorist Will Rogers said, “*Good judgment comes from experience, and a lot of that comes from bad judgment*.” An example of bad judgement was the failed attempt to eradicate wild grape (*Vitis gridiana*) in California in the early 1950s using 2,4-d herbicide as part of a program to suppress the native western grape leaf skeletonizer (*Harrisina brillians*) (Stewart et al. 1952). The suppression of the skeletonizer at public expense continued well into the 1970s, even though the skeletonizer has a spotty distribution, is highly visible, occurs primarily on grape in private gardens, and is easily managed in commercial plantings.

Regulatory agencies need to be able to assess the threat posed by an invasive species, and to be able to predict their potential geographic range and relative abundance as a measure of how favorable an area might be. *De facto* methods for assessing the potential distribution of invasive species are correlative methods that seek to circumscribe the climatic envelope of a species by identifying weather correlates based on pest incidence records, but this may fail if the species is migratory in areas that are only temporarily favorable (screwworm). However, species plasticity may enable species to accommodate weather patterns not captured by the correlates, and may lead to mistakes as to the risk posed (Davis et al. 1998). An excellent recent example of this is the invasion of wide areas of the European Palearctic region by the South American tomato pinworm (*Tuta absoluta*) (Ponti et al. 2021). Verbal and two early correlative CLIMEX (Sutherst and Maywald 1985) assessments indicated that this tropical species could not invade north temperate areas (Desneux et al. 2010, Han et al. 2019), hence, it was not identified as a quarantine pest until after its invasion of Europe, starting in 2006 when it entered Spain and then invaded colder regions of Europe (Ponti et al. 2021). Later biological studies found that the pest was moderately cold-tolerant (Kahrer et al. 2019, Li et al. 2021) and exhibited facultative diapause in response to decreasing temperatures and photoperiod (Campos et al. 2021). Incorporation of this biology in a PBDM predicted the correct invasive distribution of the pest (Ponti et al. 2021), all without reference to pest distribution records. Early, holistic, and well-parameterized PBDM assessments of this pest (and others) could have provided quarantine agencies with realistic information about its invasive potential that should have triggered quarantine measures and aided resource allocation in eradication efforts.

Weather-driven mechanistic models based on sound holistic data can be used in concert with correlative approaches to assess the prospective geographic range and abundance of invasive species. Mechanistic models of varying degrees of completeness have been used for this purpose, with PBDMs being the most common. For example, a PBDM based on European biological and ecological data predicted an extensive distribution in North America of the Palearctic grape berry moth (*Lobesia botrana*) that invaded grape (*Vitis vinifera*) in Northern California in 2009 (Gutierrez et al. 2012, Gutierrez and Ponti 2013). Fortunately, the moth was eradicated in California by early intensive detection, quarantine, and strong pheromone and chemical measures in the small initial invasion areas (Varela et al. 2010, Heit et al. 2015). The same PBDM for *Lobesia* was later used to analyze realistically the distribution and dynamics of the moth in the Palearctic region (Gutierrez et al. 2018, Ponti et al. 2018), demonstrating the transferability of the PBDM weather driven biology to other times and places. In contrast to *L. botrana*, eradication of the Australian light brown apple moth (*Epiphyas postvittana*) in California failed, but as predicted by PBDM, its range is restricted to near-coastal regions where it is a minor pest (Gutierrez et al. 2010). A good example of weather-related constraints to invasion is the failure of the semi-tropical cotton boll weevil (*Anthonomous grandis*) to invade the abundant cotton in the SW USA, more than a century and a half after it invaded the SE USA cotton. In contrast, olive fly and spotted wing drosophila (and numerous other exotic species) entered California and quickly established permanent high populations (Gutierrez et al. 2009, 2016), and no eradication efforts were conducted against them.

The eradication of the three important pests examined in this paper had varying degrees of documentation of their ecology and biology: the New World screwworm had excellent data on field ecology but poor data on its developmental biology; the Mediterranean fruit fly had moderate data on its biology and ecology; and the pink bollworm had extensive documentation on most aspects of its biology and ecology. None of the three species has effective biological control agents, and all are of sub-tropical and tropical origins. PBDM analysis of screwworm showed that it could not establish permanent populations in central Texas under extant weather, and yet massive investments were made to eradicate it there (Gutierrez and Ponti 2014, Gutierrez et al. 2019). Absent eradication, the area of permanence of the tropical screwworm northward was limited to mid-reaches of eastern Mexico, but under climate change by 2060-2065, the prospective permanent range of screwworm could increase into Texas and along the Gulf Coast and Florida.

PBDM analysis of medfly suggested it could not establish in California, and yet massive eradication campaigns occurred. Medfly’s biology is more subtropical than tropical (Gutierrez et al. 2021), explaining its establishment in some areas of the Mediterranean Basin, including northern Italy at the south end of Lake Garda. Under extant weather, only a small area of coastal southern California was predicted to be marginally suitable (FI<0.5), with medfly unlikely to establish in colder northern and central California. However, under climate change, coastal California will become more sub-tropical and become favorable for medfly.

Similarly, the cold-susceptible pink bollworm was unlikely to establish under extant weather in the Great Central Valley, where massive SIT releases were made (Gutierrez et al. 2006). However, under climate change, the prospective range of the pink bollworm would expand to all of the cotton-growing areas of California. Should reinvasion occur, the combined use of Bt cotton and SIT could be sufficient to control it.

Ricciardi et al. (2021) proffered the following priorities to deal with invasive species: (1) invasion science should strive to develop a more comprehensive framework for predicting how the behavior, abundance, and interspecific interactions of non-native species vary in relation to conditions in receiving environments and how these factors govern the ecological impacts of invasion, (2) understanding the potential synergistic effects of multiple co-occurring stressors— particularly involving climate change—on the establishment and impact of non-native species, (3) taxonomic impediments due to growing deficit in taxonomic expertise must be addressed, as new molecular technologies alone cannot adequately compensate, and (4) cooperative internationally biosecurity strategies can yield greater benefit than independent attempts by individual countries to exclude arrival and establishment of invasive species. We note that PBDMs provide a sound basis for addressing points 1 and 2 for any invasive taxa, including at the subspecies level (Gutierrez, Ponti, Levi-Mourao, et al. 2025).

Compared to the large sums spent on control and eradication efforts on these and other pest species, the costs of gathering the appropriate biological data to develop holistic weather-driven mechanistic models for them would be pitifully small and would yield considerable public benefit in evaluating their invasive potential under current weather and climate change. From the point of view of a poikilotherm species, climate change is simply another weather pattern that may alter the geographic range and relative abundance of the species, and because PBDMs model the weather-driven biology, they are time- and place-independent. Well-parameterized PBDMs can serve as an early warning system for estimating exotic pest invasiveness and for developing quarantine, containment, control, and eradication strategies. This information could forewarn regulatory agencies as to the threat posed, and at the same time, could be used to guide and assess control strategies.

